# The mechanosensitive Pkd2 channel modulates the recruitment of myosin II and actin to the cytokinetic contractile ring

**DOI:** 10.1101/2024.01.15.575753

**Authors:** Pritha Chowdhury, Debatrayee Sinha, Abhishek Poddar, Madhurya Chetluru, Qian Chen

## Abstract

Cytokinesis, the last step in cell division, separate daughter cells through the force produced by an actomyosin contractile ring assembled at the equatorial plane. In fission yeast cells, the ring helps recruit a mechanosensitive ion channel Pkd2 to the cleavage furrow, whose activation by membrane tension promotes calcium influx and daughter cell separation. However, it is unclear how the activities of Pkd2 may affect the actomyosin ring. Here, through both microscopic and genetic analyses of a hypomorphic mutant of the essential *pkd2* gene, we examine its potential role in assembling and constricting the contractile ring. The *pkd2-81KD* mutation significantly increased the number of type II myosin heavy chain Myo2 (+20%), its regulatory light chain Rlc1 (+37%) and actin (+20%) molecules in the ring, compared to the wild type. Consistent with a regulatory role of Pkd2 in the ring assembly, we identified a strong negative genetic interaction between *pkd2-81KD* and the temperature-sensitive mutant *myo2-E1*. The *pkd2-81KD myo2-E1* cells often failed to assemble a complete contractile ring. We conclude that Pkd2 modulates the recruitment of type II myosin and actin to the contractile ring, suggesting a novel calcium-dependent mechanism regulating the actin cytoskeletal structures during cytokinesis.

## Introduction

Cytokinesis is the last stage of cell division when the daughter cells separate physically through mechanical force. In most eukaryotes, this process requires an actomyosin contractile ring specifically assembled at the equatorial division plane. Several key signaling pathways regulate the assembly and constriction of this contractile ring (for a review see (Pollard and Wu, 2010)). They include those mediated by the Polo kinase, the Central Spindlin complex, and RhoA (for reviews see (Basant and Glotzer, 2018; Petronczki et al., 2008)). These pathways in turn reorganize the actin cytoskeletal structures during cytokinesis to assemble the contractile ring consisting of dozens actin-binding proteins including formin, cofilin and myosin (Balasubramanian et al., 1998; Chen and Pollard, 2011; Kovar et al., 2011; Wu and Pollard, 2005). Among them, the type II non-muscle myosin (myosin II) play a central role by driving both the assembly and disassembly of actin filaments in the contractile ring. (Balasubramanian et al., 1998; Laplante et al., 2015; Malla et al., 2022; Wu et al., 2003). Myosin II is a motor whose activities depend on its heavy chain, the essential light chain and the regulatory light chain. In fission yeast, there are two myosin II heavy chains Myo2 and Myp2, but only the former is essential for cytokinesis (Bezanilla et al., 1997). The essential light chain is Cdc4, while the regulatory light chain is Rlc1 (Le Goff et al., 2000; Naqvi et al., 2000). It remains unknown how myosin II transits from its initial role of assembling the actin filaments in the ring to its later role of disassembling these filaments as the ring constricts.

We recently found that an essential ion channel Pkd2 is required for cytokinesis of the unicellular model organism fission yeast *Schizosacchromyces pombe* (Morris et al., 2019; Palmer et al., 2005). Pkd2 belongs to the conserved family of polycystin channels. Most vertebrates possess two polycystins, PC-1 and PC-2 (for a review, see (Hu and Harris, 2020)). Mutations of human polycystins lead to a common genetic disorder, Autosomal Dominant Polycystic Kidney Disorder (ADPKD) (Consortium, 1994; Mochizuki et al., 1996). However, the cellular function of polycystins, particularly in cell division (AbouAlaiwi et al., 2011), has remained largely unknown. Fission yeast Pkd2, reconstituted in vitro, allows the passage of Ca^2+^ when the membrane is stretched by mechanical force (Poddar et al., 2022). In vivo, Pkd2 is trafficked to the plasma membrane where its distribution is cell-cycle dependent (Malla et al., 2023). Both its putative extracellular lipid-binding domain and the central nine-helix transmembrane domain are essential. In contrast, its C-terminal cytosolic tail, although dispensable, ensures its polarized distribution on the plasma membrane and selective internalization through the endocytic pathway (Malla et al., 2023). During interphase, this channel mostly localizes at the cell tips where it promotes the tip extension of fission yeast cells (Sinha et al., 2022). Upon entering cytokinesis, Pkd2 is recruited to the cleavage furrow where it stays until the end of cell separation. This Ca^2+^- permeable channel, likely activated by the membrane stretching at the cleavage furrow, promotes the separation Ca^2+^ spikes, a transient influx of Ca^2+^ during cytokinesis (Poddar et al., 2022; Poddar et al., 2021). The activity of Pkd2 is essential for separating daughter cells. To our knowledge, Pkd2 is the only calcium channel required for cytokinesis in fission yeast.

Here we tested the hypothesis that Pkd2 can regulate the assembly and constriction of the contractile ring during cytokinesis. Our hypothesis was largely based on the discovery of Pkd2 as a Ca^2+^ influx channel localized at the cleavage furrow. As an essential secondary messenger, Ca^2+^ could activate a large number of Ca^2+^-sensitive molecules including myosin regulatory light chains, calmodulin, calcineurin and Ca^2+^-dependent kinases. Many of them play a crucial role in regulating the actin cytoskeletal structures. In this study, we took advantage of a hypomorphic mutant of the essential *pkd2* gene. This mutation *pkd2-81KD* replaces the endogenous promoter with an inducible 81nmt1 promoter, reducing the expression of *pkd2* gene by more than 70% under the suppressing condition (Morris et al., 2019). One of the surprising phenotypes of this mutant is that it accelerates the cleavage furrow ingression, even though the mutant daughter cells often fail to separate (Morris et al., 2019). Here, using quantitative fluorescence microscopy, we compared the number of type II myosin heavy chain Myo2, the regulatory light chain Rlc1 and actin molecules in the contractile ring of *pkd2-81KD* to the wild type. In addition to the microscopic analyses, we examined the genetic interaction between the *pkd2* mutants, including both *pkd2-81KD* and the new temperature-sensitive mutant *pkd2-B42* (Sinha et al., 2022), and the *myo2-E1* mutant. Overall, our results would demonstrate that Pkd2 modulates the recruitment of both myosin II and actin to the contractile ring, suggesting that this force-activated channel may regulate the assembly of actin cytoskeletal structures during cytokinesis through Ca^2+^.

## Materials and methods

### Cell culture and Yeast genetics

Yeast cell culture and genetics were carried out according to the standard protocols (Moreno et al., 1991). Unless specified, YE5S was used in all experiments. For genetic crosses, tetrads were dissected using Sporeplay+ dissection microscope (Singer, England). Yeast strains used in this study are listed in Table 1. For ten-fold dilution series of yeast, overnight cultures were diluted and inoculated for additional 5 hours before being spotted onto YE5S agar plates. The plates were incubated for 2-3 days in respective temperatures before being scanned using a photo scanner (Epson).

**Table 1.** List of yeast strains used in this study.

| **Name** | **Genotype** | **Source** |
| --- | --- | --- |
| FY527 | *h- leu1-32 ura4-D18 his3-D1 ade6-M216* | Lab stock |
| JW766 | *h+ kanMX6-Pmyo2-GFP-myo2 ade6-M210 leu1-32 ura4-D18* | Lab stock |
| JW1341 | *h- rlc1-tdTomato-natMX6 ade6-M210 leu1-32 ura4-D18* | Lab stock |
| QC-Y601 | *h+ leu2::kanMX-Pcof1-mEGFP-Lifeact rlc1-tdTomato-Nat* | Lab stock |
| QC-Y799 | *h+ myo2-E1 rlc1-tdTomato-NatMX6 sad1-mGFP-KanMX6* | This study |
| QC-Y813 | *h- kanMX6-81xnmt1-pkd2 rlc1-tdTomato-NatMX6* | Lab stock |
| QC-Y817 | *h+ kanMX6-81xnmt1-pkd2 leu1-32 ura4-D18 his3-D1 ade6-M210* | Lab stock |
| QC-Y839 | *h- myo2-E1* |  |
| QC-Y840 | *h+ myo2-E1 kanMX6-81xnmt1-Pkd2* | This study |
| QC-Y1033 | *h? myo2-E1 kanMX6-81xnmt1-pkd2 rlc1-tdTomato-natMX6* | This study |
| QC-Y1112 | *h? kanMX6-Pmyo2-GFP-myo2 ade6-M210 leu1-32 ura4-D18 kanMX6-81xnmt1-pkd2* | This study |
| QC-Y1156 | *h? leu2::kanMX-Pcof1-mEGFP-Lifeact rlc1-tdTomato-Nat KanMX6-P81nmt1-pkd2 leu1-32 ura4-D18 his3-D1* | Lab stock |
| QC-Y1752 | *h? pkd2::pkd2-B42-ura4+-hist5+ leu1-32 myo2-E1* | This study |

### Microscopy

For microscopy, fission yeast cells were first inoculated in YE5S liquid media for two days at 25°C before being harvested through centrifugation at 1500g for 1 min during exponential phase at a density between 5x10^6^ cells/ml and 1.0x10^7^ cells/ml. The cells were resuspended in 50 µl of YE5S and 6 µl of the resuspended cells were spotted onto a gelatin pad (25% gelatin in YE5S) on a glass slide. The cells were sealed under a coverslip (#1.5) with VALAP (1:1:1 mixture of Vaseline, lanolin and paraffin) before imaging.

Live microscopy was carried out on a spinning disk confocal microscope equipped with EM-CCD camera. The Olympus IX71 microscope was equipped with three objective lenses of 60x (NA= 1.40, oil) and 100x (NA= 1.40, oil), a motorized Piezo Z Top plane (ASI, USA) and a confocal spinning disk unit (CSU-X1, Yokogawa, Japan). The images were captured on an Ixon- 897 EMCCD camera (Andor). Solid state lasers of 488 and 561 nm were used to excite green (GFP) and red fluorescence (tdTomato) proteins respectively. Unless specified, the cells were maged with Z-series of 15 slices at a step size of 0.5 µm.

To stain the cell wall with calcofluor, exponentially growing cell culture was fixed with 4% paraformaldehyde, prepared from 16% fresh stock solution (Electron Microscopy Science, USA). The fixed cells were washed with TEMK buffer (50 mM Tris-HCL pH 7.4, 1 mM MgCl2, 50 mM KCl,1 mm EGTA, pH 8.0). The fixed cells were then stained with 1 µg/ml calcofluor, prepared with the stock solution (1 mg/ml, Sigma Aldrich) on a rocking platform for 10 minutes at the room temperature in the dark. The stained cells were pelleted and resuspended in 50 µl TEMK, 6 µl of which was spotted on a glass slide directly and sealed under a coverslip (#1.5) with VALAP. The stained samples were imaged with an Olympus IX81 microscope equipped with 60x objective lens (NA=1.42), a LED lamp for bright-field microscopy, and a mercury lamp for fluorescence microscopy. The micrographs were captured on an ORCA C- 11440 digital camera (Hamamatsu, Japan), operated with the CellSens software (Olympus).

### Western Blots

For probing the expression level of GFP-Myo2, 50ml of exponentially growing yeast cells were harvested by centrifugation at 1500g for 5 min. We washed the cells with 1ml of sterile water once. The pelleted cells were re-suspended with 300µl of lysis buffer (50mM Tris-HCl pH 7.5, 100 mM KCl, 3 mM MgCl2, 1 mM EDTA, 1 mM DTT, 0.1% Triton X-100 and protease inhibitors (Halt protease inhibitor cocktail; #1862209; Thermo-Fisher)). The cells were mixed with 300 mg of glass beads (0.5mm) before being mechanically homogenized by a bead beater (BeadBug, Benchmark Scientific) for five cycles of 1min breaking interrupted by 1min incubation on ice. The lysed cell was immediately mixed with pre-heated 5x sample buffer (250mM Tris-HCL pH 6.8, 50% glycerol, 25% β-mercaptoethanol, 15% SDS, 0.025% Bromophenol Blue) before being heated for 10 minutes at 100°C. After centrifugation at top speed for 1 minute, the supernatant was collected for SDS-PAGE gel electrophoresis (Mini-PROTEIN TGX 10% precast, BioRad #4561033). The gels were either stained with Coomassie blue or transferred to PVDF membrane (Amersham #10600023) for immunoblots. The membrane was blotted with anti-GFP monoclonal primary antibody (mouse monoclonal, Sigma-Roche #11814460001) at 4°C overnight, followed by horseradish peroxidase-conjugated Goat-anti-mouse second antibodies (1:10000; BioRad # 1721011) for 1hr at the room temperature.

Blots were developed with chemifluorescent reagents (Pierce ECL Western Blotting Substrate Thermo Scientific #32209). The blots were quantified through ImageJ.

### Image analysis

We used ImageJ (NIH), its plug-ins and macros to process all the micrographs. For quantitative analysis, the time-lapse series were first corrected for X-Y drifting using the plugin StackReg (Thevenaz et al., 1998). Average intensity projections of Z-series were used for quantification. The localization of GFP-Myo2 and Rlc1-tdTomato at the contractile ring was quantified by measuring the fluorescence intensities in a 1 µm by 4 µm rectangle centered on the equatorial division plane. The measurements were corrected for the background fluorescence, measured in a 0.2 µm by 4 µm rectangle adjoined to the equatorial division plane. To measure fluorescence intensities of GFP-Lifeact in the contractile ring, we used the fluorescence of Rlc1-tdTomato to manually segment the contractile ring. On average the contractile ring of a wild type covered 1.38 µm^2^. The mean fluorescence of the GFP-Lifeact was multiplied by the average area of wild type ring. To determine the start of the contractile ring constriction, we analyzed the fluorescence kymographs constructed from the time-lapse series of the contractile ring.

For quantification of immunoblots, the band intensities were measured by polygon tools of ImageJ and the measurements were corrected for background fluorescence.

All the figures were prepared with Canvas X (ACD Systems) and the plots were prepared using Origin 2021 (OriginLab, USA).

## Results

### Pkd2 modulates the recruitment of Myo2, Rlc1, and actin to the contractile ring

We first determined how Pkd2, as a Ca^2+^ influx channel localized at the cleavage furrow, regulates the localization of the myosin II heavy chain Myo2 to the contractile ring. To measure the relative number of Myo2 molecules in the *pkd2-81KD* mutant cells, we compared the fluorescence intensities of the N-terminally tagged GFP-Myo2, expressed from its endogenous locus, of the mutant to that of the wild type using quantitative microscopy (Malla et al., 2022) (Fig. 1A). The intracellular concentration of GFP-Myo2, measured by either the fluorescence intensities (Fig. 1B) or the immunoblots (Fig. 1C) was unchanged in the mutant cells. During cytokinesis, similar to the wild type cells, the *pkd2* mutant cells recruited GFP-Myo2 to a broad band of cytokinesis nodes before condensing the nodes into a complete contractile ring (Fig. 1D). However, the contractile ring of the *pkd2* mutant cells matured and constricted more quickly than that of the wild type cells (Fig. 1D), as reported previously (Morris et al., 2019). The fluorescence intensity of GFP-Myo2 in the contractile ring of the *pkd2* mutant cells increased more quickly and peaked sooner than the wild type (Fig. 1E). As a result, the fluorescence intensity of GFP-Myo2 in a mature ring, referred to the ring just before the cleavage furrow ingression, of the mutant increased by 18% (P<0.01) (Fig. 1F). Thus, Pkd2 modulates the recruitment of the myosin II heavy chain Myo2 to the contractile ring during cytokinesis.

**Figure 1:**
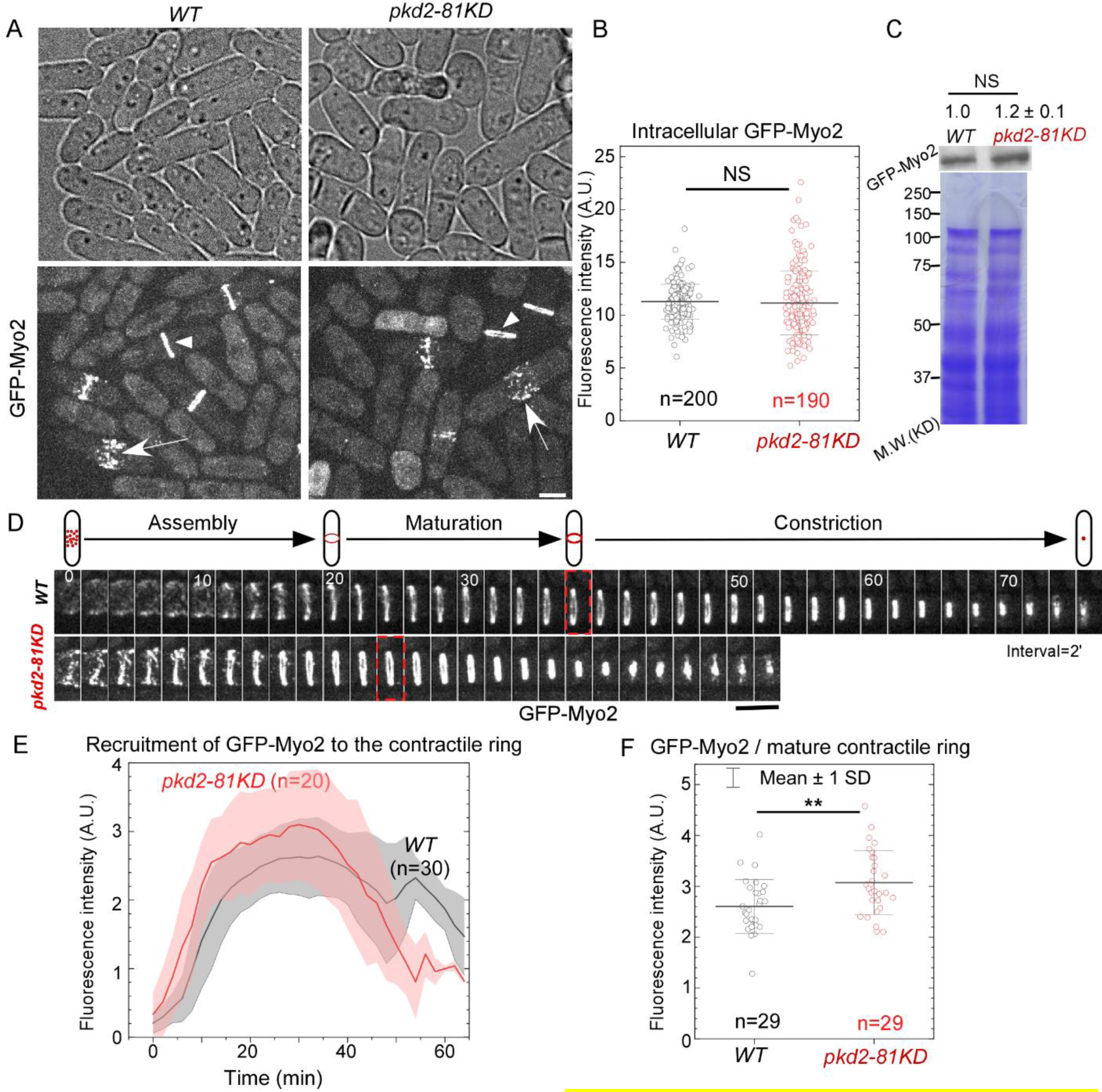
The *pkd2-81KD* mutant increased the recruitment of myosin II heavy chain Myo2 to the contractile ring. (A) Micrographs of wild type and *pkd2-81KD* cells expressing GFP- Myo2.Arrow: nodes of contractile ring; Arrowhead: the contractile ring. (**B)** Dot plot of the average intracellular fluorescence intensity of GFP-Myo2. **(C)** Expression of GFP-Myo2 in the *pkd2* mutant cells. Top: anti-GFP blot of the lysate from wild type and *pkd2-81KD* cells expressing GFP-Myo2. Bottom: Coomassie blue stained SDS-PAGE gel of respective lysates. Representative data from three biological repeats are shown. **(D)** Top: a diagram of the process of contractile ring assembly, maturation and constriction in a fission yeast cell. Bottom: time series of the equatorial plane of dividing cells expressing GFP-Myo2. Dashed box: start of the contractile ring constriction. Number: time in minutes. **(E)** Time course of the fluorescence intensity of GFP-Myo2 at the equatorial plane after the start of contractile ring assembly (time zero). Cloud: standard deviations. **(F)** Dot plot of the fluorescence intensity of GFP-Myo2 in a mature contractile ring just before it starts to constrict. *: P<0.05. **: P<0.01. ***: P<0.001. N.S.: not significant. Statistics were calculated using two-tailed Student’s t tests. All the data are pooled from at least two independent biological repeats. Scale bar = 5 µm.

Next, we examined whether Pkd2 helps recruit Rlc1, the regulatory light chain of myosin II (Naqvi et al., 2000), to the contractile ring. Rlc1 possesses four EF-hands which bind Ca^2+^. It is a potential target of the Ca^2+^ signaling pathway regulated by Pkd2. As above, to measure the relative number of Rlc1 molecules in the *pkd2* mutant cells, we compared the fluorescence intensity of Rlc1, tagged endogenously with the fluorescence protein tdTomato (Rlc1-tdTomato), in *pkd2-81KD* cells to that in the wild type cells (Fig. 2A). Interestingly, we found a significant increase (26%, P<0.001) in the intracellular fluorescence intensities of Rlc1-tdTomato of the *pkd2* mutant (Fig. 2B). During cytokinesis, Rlc1-tdTomato was recruited to a broad band of nodes in the *pkd2* mutant cells before these nodes condensed into the contractile ring (Fig. 2C).

**Figure 2:**
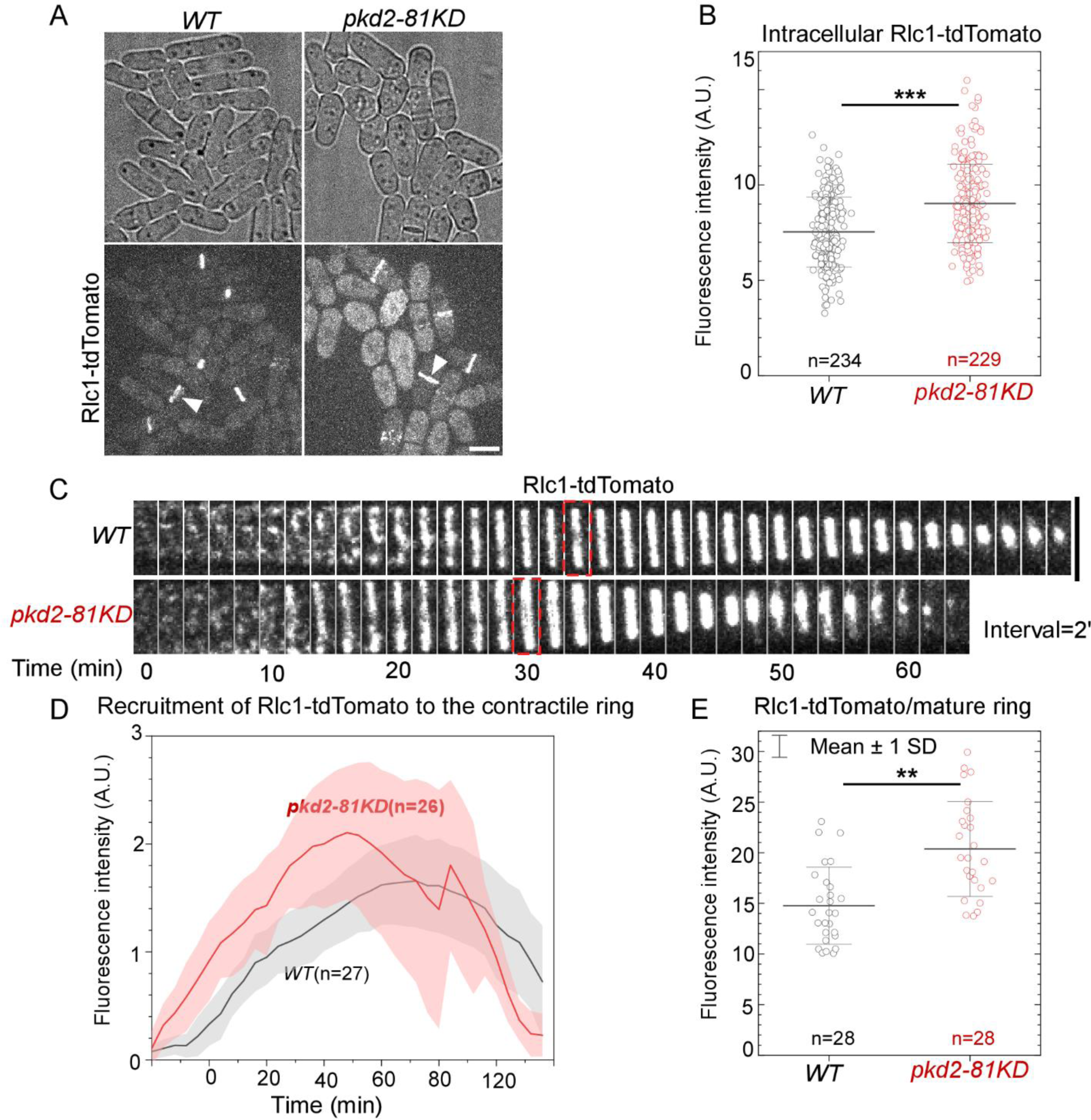
The *pkd2-81KD* mutant increased the recruitment of myosin II regulatory light chain Rlc1 to the contractile ring. (A) Micrographs of *wild type* and *pkd2-81KD* cells expressing Rlc1-tdTomato. Arrowhead: the contractile ring. **(B)** Dot plot of the intracellular fluorescence intensity of Rlc1-tdTomato. **(C)** Time series of the equatorial plane of dividing cells expressing Rlc1-tdTomato. Dashed box: start of the contractile ring constriction. Number indicates time in minutes. **(D)** Time course of the fluorescence intensity of Rlc1-tdTomato in the equatorial plane. Cloud: standard deviations. **(E)** Dot plot of the fluorescence intensity of Rlc1-tdTomato in a mature contractile ring. *: P<0.05. **: P<0.01. ***: P<0.001. N.S.: not significant. Statistics were calculated using two-tailed Student’s t-tests. All the data are pooled from at least two independent biological repeats. Scale bar = 5 µm.

However, the *pkd2-81KD* mutant resulted in the fluorescence of Rlc1-tdTomato in the ring increasing much more quickly and peaking at least 10 mins sooner (Fig. 2D). Similar to Myo2, the average fluorescence intensity of Rlc1-tdTomato in a mature ring of the mutant was 37% higher than that of the wild type (P<0.001) (Fig. 2E). Therefore, in addition to the myosin II heavy chain, Pkd2 also modulates the recruitment of the regulatory light chain to the cytokinetic contractile ring.

Lastly, we determined whether Pkd2 has a role in the assembly of actin filaments in the cytokinetic contractile ring. We employed GFP-Lifeact as the fluorescence reporter (Malla et al., 2022) to measure the number of actin molecules in the ring. The endogenously expressed GFP- Lifeact was driven by a promoter of intermediate strength (Pcof1) to minimize the potential of interfering with the actin cytoskeleton by this probe (Courtemanche et al., 2016). The *pkd2* mutation substantially altered all the actin cytoskeletal structures including endocytic actin patches, actin cables and the contractile ring (Fig. 3A). In particular, the number of actin patches increased significantly in the *pkd2-81KD* mutant cells, compared to the wild type. Compared to the wild type cells in which the actin patches concentrated at the cell tips, more patches of the mutant spread out in the middle portion of a cell (Fig. 3A). Interestingly, the intracellular concentration of GFP-Lifeact in the mutant cells increased dramatically by almost 90% (P<0.001) (Fig. 3B). During cytokinesis, the assembly and disassembly of actin filaments, labeled by GFP-Lifeact, proceeded normally in the contractile ring of the mutant cells (Fig. 3C). The fluorescence intensities of GFP-Lifeact in the ring gradually increased before the ring constriction. They decreased as the ring constricted. We again found that the fluorescence of GFP-Lifeact in the contractile ring of *pkd2-81KD* cells was 21% higher than the wild type cells (P<0.001) (Fig. 3D), similar to myosin II. Based on these results, we concluded that in addition to myosin II, Pkd2 also modulates the assembly of actin filaments in the cytokinetic contractile ring.

**Figure 3:**
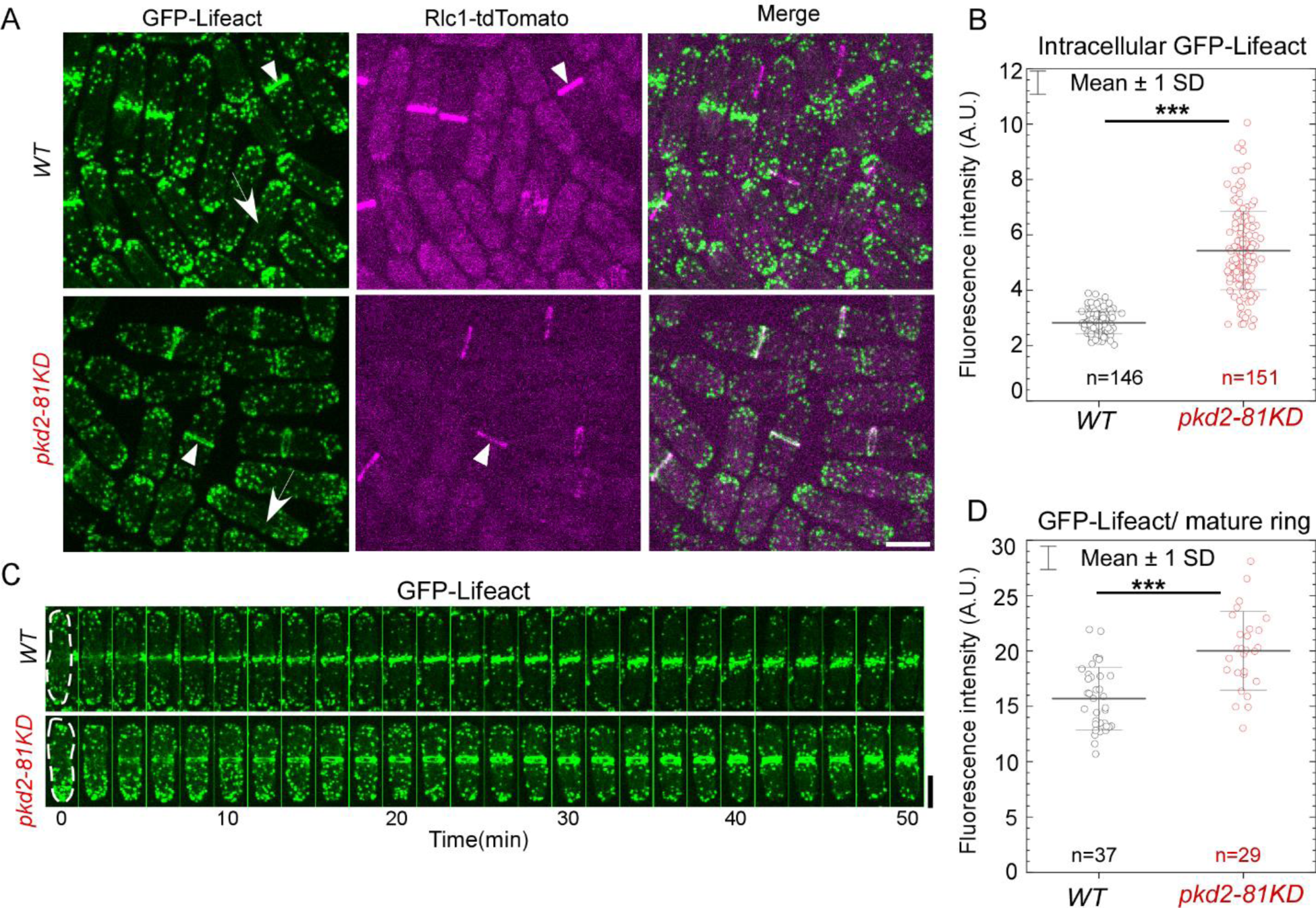
The *pkd2*-*81KD* mutant increased the assembly of actin filaments in the contractile ring. (A) Micrographs of the *wild type* (top) and *pkd2-81KD* (bottom) cells co-expressing GFP-Lifeact (green) and Rlc1-tdTomato (magenta). Arrow: the actin cables; Arrowhead: the contractile ring. **(B)** Dot plot of the intracellular fluorescence intensity of GFP-Lifeact. **(C)** Time series of a *wild type* and a *pkd2-81KD* cell expressing GFP-Lifeact. Number: Time in minutes from the start of the contractile ring assembly. **(D)** Dot plot of the fluorescence intensity of GFP-Lifeact in a mature contractile ring. ***: P<0.001. N.S.: not significant. Statistics were calculated using two-tailed Student’s t tests. All the data are pooled from at least two independent biological repeats. Bar=5µm.

### Both *pkd2* and *myo2* are essential for the contractile ring assembly in cytokinesis

To further understand the relationship between *pkd2* and *myo2*, we examined their interactions through genetic studies of *pkd2* and *myo2* mutants. First, we confirmed the previously discovered negative genetic interaction between *pkd2-81KD* and a temperature-sensitive mutant *myo2-E1* (Fig. 4A) (Morris et al., 2019). Now, we similarly found a negative genetic interaction between the temperature-sensitive mutant *pkd2-B42* (Sinha et al., 2022) and *myo2-E1* (Fig. 4A). Even at the permissive temperature of 25°C, the double mutant *pkd2-B42 myo2-E1* grew much more slowly than either *pkd2-B42* or *myo2-E1* (Fig. 4A). Next, we compared the morphogenesis of *pkd2-81KD, myo2-E1*, and *pkd2-81KD myo2-E1* cells with each other. The morphology of *pkd2- 81KD myo2-E1* cells was similar to that of the *pkd2-81KD* cells (Fig. 4B). Both were rounder than the wild type cells. Both contained a small fraction of “deflated” cells, which shrank temporarily even in the rich YE medium (Fig. 4B). However, the *pkd2-81KD myo2-E1* mutant stood out for its large fraction of lysed cells (11±1.7%, average ± standard deviation) even at the permissive temperature of 25°C, far more than either the *pkd2-81KD* or *myo2-E1* mutant (Fig.4B). A much higher fraction of the double mutant (74%) contained septum than either of the single mutants (Fig. 4C and 4D). The septum of *pkd2-81KD myo2-E1* cells often appeared to be wavy, a rare phenotype among either of the single mutants (Fig. 4C). Much more double mutant cells were multi-septated (21±3%) than either *pkd2-81KD* or *myo2-E1* cells (Fig. 4D). Based on these genetic analyses, we concluded that *pkd2* and *myo2* have a synergistic relationship in cytokinesis.

**Figure 4:**
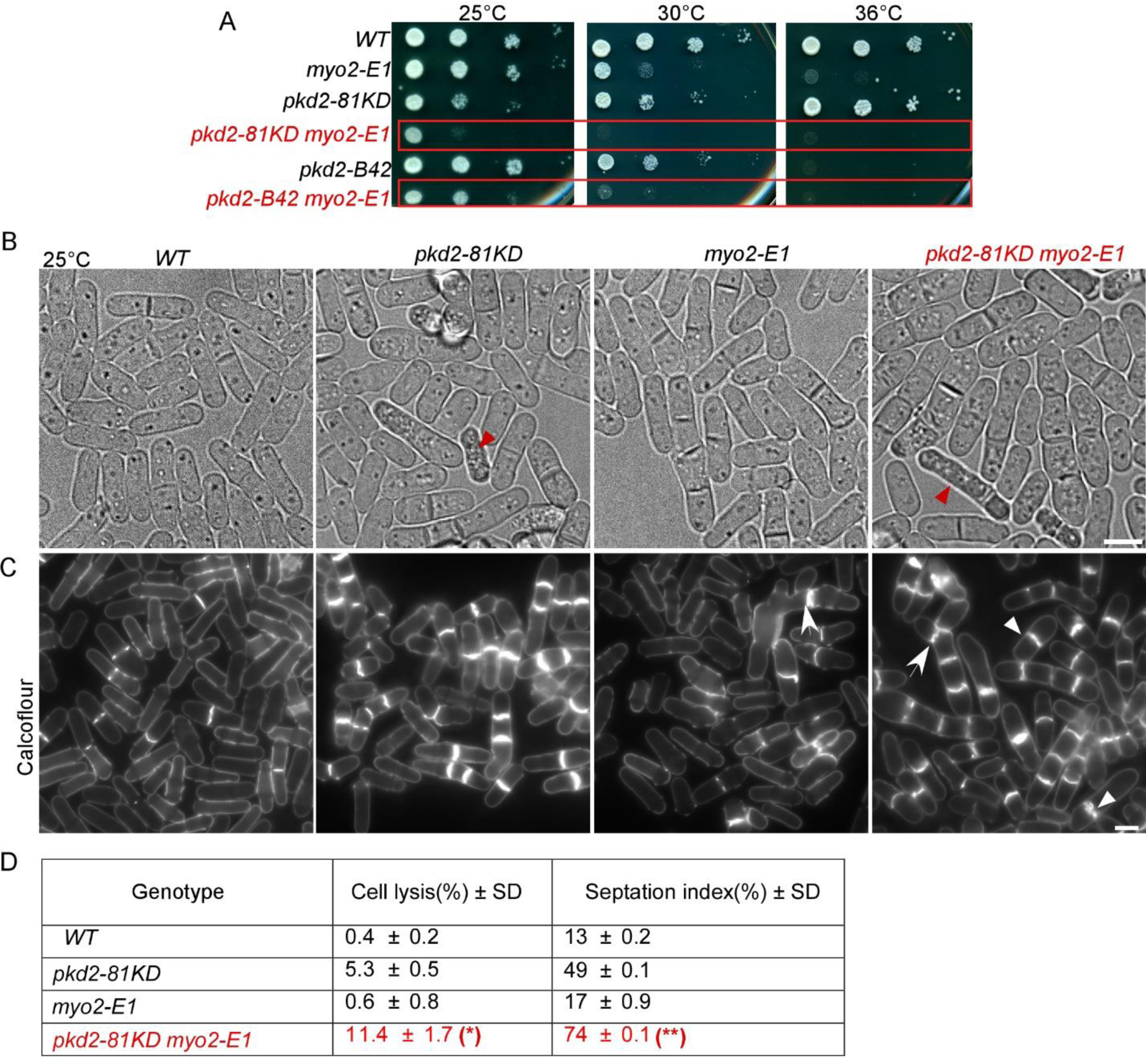
Negative genetic interactions between *pkd2* and *myo2* mutants. (A) Ten-fold dilution series of yeast cells at the indicated temperatures. **(B)** Bright-field micrographs of live cells at 25°C. Red arrowhead: lysed cells. **(C)** Micrographs of calcofluor-stained fixed cells at 25°C. Arrow: abnormal septum. Arrowhead: thick septum (n>500). **(D)** A table summarizing the morphological defects of the *pkd2-81KD myo2-E1* mutant cells. All the data are pooled from at least two independent biological repeats. ***: P<0.001. Statistics were calculated using two- tailed Student’s t tests. Scale bar = 5 µm.

To determine how Pkd2 and Myo2 work synergistically in cytokinesis, we measured the contractile ring assembly and constriction in the *pkd2-81KD myo2-E1* cells. We used Rlc1- tdTomato as the marker for the contractile ring in these double mutant cells. We immediately noticed, even at the permissive temperature of 25°C, that the number of the contractile rings appeared to be far lower in *pkd2-81KD myo2-E1* cells than *myo2-E1* cells (Fig. 5A).

**Figure 5:**
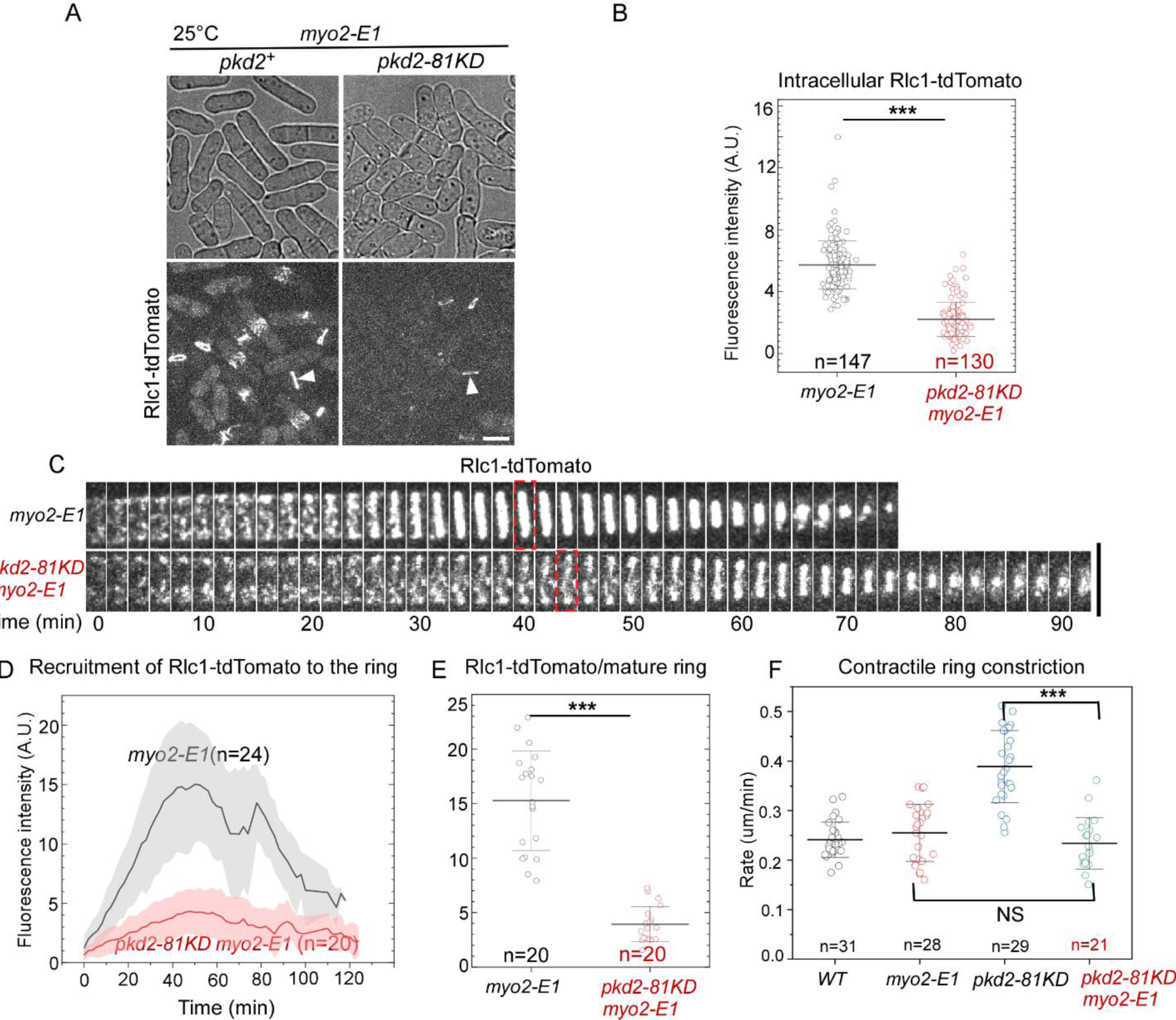
The *pkd2-81KD myo2-E1* mutant failed to assemble a complete contractile ring at the permissive temperature. (A) Micrographs of live cells expressing Rlc1-tdTomato. Arrowhead: the contractile ring. **(B)** Dot plot of intracellular fluorescence intensity of Rlc1- tdTomato. **(C)** Time-lapse series of the equatorial plane of a *myo2-E1* and a *pkd2-81KD myo2- E1* cell expressing Rlc1-tdTomato. Dashed box: start of contractile ring constriction. Number: time in minutes. **(D)** Time course of the fluorescence intensity of Rlc1-tdTomato at the equatorial plane. **(E)** Dot plot of average fluorescence intensity of Rlc1-tdTomato in a mature contractile ring before it starts to constrict. **(F)** Dot plot of the rate of the contractile ring constriction. All the experiments were carried out at the permissive temperature of 25°C. ***: P<0.001. N.S.: not significant. Statistics were calculated using two-tailed Student’s t tests. All the data are pooled from at least two independent biological repeats. Scale bar = 5 µm.

Quantitative measurement revealed that the intracellular fluorescence of Rlc1-tdTomato in the *pkd2-81KD myo2-E1* cells was 70% lower than that of *myo2-E1* (P<0.001) (Fig. 5B). Although the double mutant cells condensed the cytokinetic nodes into a contractile ring, most of these rings were incomplete with many gaps before the constriction ensued (n=26) (Fig. 5C). The fluorescence of Rlc1-tdTomato in the ring increased very slowly in the *pkd2-81KD myo2-E1* mutant cells, but it peaked at the same time as the wild type cells (Fig. 5D). As a result, the average fluorescence of Rlc1-tdTomato in a mature ring decreased by 74% in the double mutant, compared to *myo2-E1* (P<0.001) (Fig. 5E). Surprisingly, these rings constricted at a rate comparable to those of either wild type or *myo2-E1* cells at 25°C, but much more slowly than *pkd2-81KD* cells (Fig. 5F). We concluded that both the myosin II heavy chain Myo2 and Pkd2 work synergistically to recruit the myosin regulatory light chain Rlc1 to the contractile ring.

## Discussion

In this study, we investigated a potential role of the mechanosensitive ion channel Pkd2 in the assembly and constriction of the actomyosin contractile ring during cytokinesis. We found that Pkd2 modulates the recruitment of myosin II heavy chain, its regulatory light chain and actin to the contractile ring. The regulatory role of Pkd2 is synergistic with that of myosin II in promoting the assembly of a complete contractile ring.

The modulating role of Pkd2 in recruiting myosin II and actin to the contractile ring, although surprising, is consistent with the cytokinesis defects of the *pkd2* mutant cells. Our earlier works have demonstrated that the cleavage furrow ingressed almost 50% more quickly in the *pkd2* mutant cells compared to the wild type (Morris et al., 2019). This is partially due to the reduced turgor pressure of the mutant cells (Sinha et al., 2022). Here, we found another factor that may contribute to the faster ingression of the cleavage furrow in the *pkd2* mutant cells. The contractile ring of the mutant contains significantly more myosin II and actin molecules than the wild type. The myosin II heavy chain Myo2 increases proportionally to actin filaments in the ring of the *pkd2-81KD* mutant cells, but it is substantially less than the increase of the regulatory light chain Rlc1 in the ring. Although the increase of the heavy chain by itself has little effect on the constriction of the contractile ring (Stark et al., 2010), the increased densities of both Myo2 and the regulatory light chain Rlc1 as well as actin filaments may. These three molecules together shall allow the ring to generate additional mechanical force during cytokinesis of the *pkd2-81KD* cells. Combining this increased compression force with the reduced resistance from the turgor pressure in the *pkd2* mutant cells, it is not surprising that the cleavage furrow ingresses much more quickly in *pkd2-81KD* mutant cells.

The modulating effect of Pkd2 in the recruitment of myosin II and actin to the contractile ring is most likely Ca^2+^-dependent. Although the role of Ca^2+^ in fission yeast cytokinesis remains unclear, this secondary messenger plays a critical role in cytokinesis of animal cells. In these cells, Ca^2+^-dependent myosin light chain kinase (MLCK) phosphorylates the myosin II regulatory light chain, triggering the contractile ring constriction (Craig et al., 1983). However, such a mechanism of action by Ca^2+^ is not evolutionarily conserved. Unlike animal cells, fission yeast does not possess a homologue of MLCK. Instead, Rlc1 is phosphorylated by the P21- acivated kinases Pak1 and Pak2 during cytokinesis (Loo and Balasubramanian, 2008; Prieto-Ruiz et al., 2023). Although such modification of Rlc1 is not essential for cytokinesis under normal growth condition (McCollum et al., 1999), it does become necessary when the availability of actin is limited under stress condition (Prieto-Ruiz et al., 2023). So far, there is no evidence that either Pak1 or Pak2 depend on Ca^2+^/calmodulin for their kinase activities. It remains thus in doubt whether these two kinases could be the downstream targets of Pkd2.

Alternatively, the phosphorylation of Rlc1 may be regulated through a balance of phosphorylation by Pak1/2 and dephosphorylation by the calcium/calmodulin-activated phosphatase calcineurin Ppb1 (Martin-Garcia et al., 2018; Yoshida et al., 1994). Further work will be needed to illustrate how Pkd2 modulates the recruitment of myosin II through Ca^2+^.

In addition to direct modulation of the recruitment of myosin and actin molecules, Pkd2 may regulate the transcription of these genes. This mechanism may offer an alternative explanation to why mutations of *pkd2* increase the intracellular concentration of Rlc1 and GFP-Lifeact. Specifically, the significant increase of intracellular concentration of GFP-Lifeact in the *pkd2* mutant cells suggests that the expression of this actin probe may be affected by the *pkd2* mutation. Similarly, we have observed elevated transcription of GFP in the temperature-sensitive *pkd2-B42* cells (Sinha et al., 2022). The molecular mechanism of such transcription regulation by Pkd2 remains unclear. One plausible scenario is that the cytoplasmic tail of Pkd2 may translocate to nucleus to directly regulate transcription as its human homologues PC-1 does (Chauvet et al., 2004). However, we have not observed even partial localization of Pkd2 in the nucleus so far.

Alternatively, Pkd2 could indirectly regulates transcription through calcineurin. The Ca^2+^/calmodulin-dependent phosphatase Ppb1 promotes the nuclear translocation and activation of the transcription activator Prz1 (Hirayama et al., 2003). Future work will determine the potential mechanism by which Pkd2 regulates gene transcription.

Considering the modulating effects of Pkd2 in the recruitment of Myo2 to the contractile ring, the negative genetic interaction between the *pkd2* and *myo2* mutants is surprising. Even more surprising is our discovery that the abundance of Rlc1 in the contractile ring decreased dramatically in the contractile ring of the *myo2 pkd2* double mutant cells. This is despite the fact that *pkd2-81KD* mutation increases the recruitment of regulatory light chain of Myosin II to the contractile ring. A possible explanation is that both Pkd2 and Myo2 promote the stability of the contractile ring. Pkd2 may stabilize the contractile ring through modulating the assembly of actin and myosin II. In contrast, the myosin heavy chain Myo2 promotes the contractile ring stability through its motor activity, which is compromised in the *myo2-E1* mutant (Balasubramanian et al., 1998). Our discovery suggests that both Pkd2 and Myo2 stabilize the contractile ring although they do so through disparate mechanisms.

Overall, this study demonstrated that the force-sensitive ion channel Pkd2 regulates the assembly of the actomyosin contractile ring through modulating the recruitment of myosin II and actin to the ring. Our findings suggest that Ca^2+^ and Ca^2+^-binding proteins may play a critical role in reorganizing the actin cytoskeletal structures during fission yeast cytokinesis despite a lack of the canonical MLCK-dependent pathway.

## Author contributions

Conceptualization, QC.; Methodology, QC, PC and DS; Software, QC; Validation, QC, PC, DS, AP and MD; Formal Analysis, QC and PC; Investigation, QC, PC, DS, AP and MD; Resources, QC, PC and DS; Data Curation, QC, PC and DS; Writing – Original Draft Preparation, QC and PC; Writing – Review & Editing, QC and PC; Visualization, QC and PC; Supervision, QC; Project Administration, QC; Funding Acquisition, QC.

## Acknowledgements

We would like to thank the Chen lab members at The University of Toledo for technical assistance. This work has been supported by the National Institutes of Health grant R01GM144652 to QC and the National Science Foundation grant 2144701 to QC. The content is solely the responsibility of the authors and does not necessarily represent the official views of the National Institutes of Health. The authors declare no conflicts of interests.

## References

1. Morris, Z., D. Sinha, A. Poddar, B. Morris, and Q. Chen. 2019. Fission yeast TRP channel Pkd2p localizes to the cleavage furrow and regulates cell separation during cytokinesis. Molecular biology of the cell. 30:1791–1804.

2. Courtemanche, N., T.D. Pollard, and Q. Chen. 2016. Avoiding artefacts when counting polymerized actin in live cells with LifeAct fused to fluorescent proteins. Nature cell biology. 18:676–683.

3. Malla, M., T.D. Pollard, and Q. Chen. 2022. Counting actin in contractile rings reveals novel contributions of cofilin and type II myosins to fission yeast cytokinesis. Molecular biology of the cell. 33:ar51.

4. AbouAlaiwi, W.A., S. Ratnam, R.L. Booth, J.V. Shah, and S.M. Nauli. 2011. Endothelial cells from humans and mice with polycystic kidney disease are characterized by polyploidy and chromosome segregation defects through survivin down-regulation. Human molecular genetics. 20:354–367.

5. Balasubramanian, M.K., D. McCollum, L. Chang, K.C. Wong, N.I. Naqvi, X. He, S. Sazer, and K.L. Gould. 1998. Isolation and characterization of new fission yeast cytokinesis mutants. Genetics. 149:1265–1275.

6. Basant, A., and M. Glotzer. 2018. Spatiotemporal Regulation of RhoA during Cytokinesis. Curr Biol. 28:R570–R580.

7. Bezanilla, M., S.L. Forsburg, and T.D. Pollard. 1997. Identification of a second myosin-II in Schizosaccharomyces pombe: Myp2p is conditionally required for cytokinesis. Molecular biology of the cell. 8:2693–2705.

8. Chauvet, V., X. Tian, H. Husson, D.H. Grimm, T. Wang, T. Hiesberger, P. Igarashi, A.M. Bennett, O. Ibraghimov-Beskrovnaya, S. Somlo, and M.J. Caplan. 2004. Mechanical stimuli induce cleavage and nuclear translocation of the polycystin-1 C terminus. The Journal of clinical investigation. 114:1433–1443.

9. Chen, Q., and T.D. Pollard. 2011. Actin filament severing by cofilin is more important for assembly than constriction of the cytokinetic contractile ring. The Journal of cell biology. 195:485–498.

10. Consortium, E.P.K.D. 1994. The polycystic kidney disease 1 gene encodes a 14 kb transcript and lies within a duplicated region on chromosome 16. The European Polycystic Kidney Disease Consortium. Cell. 77:881–894.

11. Courtemanche, N., T.D. Pollard, and Q. Chen. 2016. Avoiding artefacts when counting polymerized actin in live cells with LifeAct fused to fluorescent proteins. Nature cell biology. 18:676–683.

12. Craig, R., R. Smith, and J. Kendrick-Jones. 1983. Light-chain phosphorylation controls the conformation of vertebrate non-muscle and smooth muscle myosin molecules. Nature. 302:436–439.

13. Hirayama, S., R. Sugiura, Y. Lu, T. Maeda, K. Kawagishi, M. Yokoyama, H. Tohda, Y. Giga-Hama, H. Shuntoh, and T. Kuno. 2003. Zinc finger protein Prz1 regulates Ca2+ but not Cl- homeostasis in fission yeast. Identification of distinct branches of calcineurin signaling pathway in fission yeast. The Journal of biological chemistry. 278:18078–18084.

14. Hu, J., and P.C. Harris. 2020. Regulation of polycystin expression, maturation and trafficking. Cellular signalling. 72:109630.

15. Kovar, D.R., V. Sirotkin, and M. Lord. 2011. Three’s company: the fission yeast actin cytoskeleton. Trends in cell biology. 21:177–187.

16. Laplante, C., J. Berro, E. Karatekin, A. Hernandez-Leyva, R. Lee, and T.D. Pollard. 2015. Three myosins contribute uniquely to the assembly and constriction of the fission yeast cytokinetic contractile ring. Curr Biol. 25:1955–1965.

17. Le Goff, X., F. Motegi, E. Salimova, I. Mabuchi, and V. Simanis. 2000. The S. pombe rlc1 gene encodes a putative myosin regulatory light chain that binds the type II myosins myo3p and myo2p. Journal of cell science. 113 Pt 23:4157–4163.

18. Loo, T.H., and M. Balasubramanian. 2008. Schizosaccharomyces pombe Pak-related protein, Pak1p/Orb2p, phosphorylates myosin regulatory light chain to inhibit cytokinesis. The Journal of cell biology. 183:785–793.

19. Malla, M., T.D. Pollard, and Q. Chen. 2022. Counting actin in contractile rings reveals novel contributions of cofilin and type II myosins to fission yeast cytokinesis. Molecular biology of the cell. 33:ar51.

20. Malla, M., D. Sinha, P. Chowdhury, B.T. Bisesi, and Q. Chen. 2023. The cytoplasmic tail of the mechanosensitive channel Pkd2 regulates its internalization and clustering in eisosomes. Journal of cell science. 136.

21. Martin-Garcia, R., V. Arribas, P.M. Coll, M. Pinar, R.A. Viana, S.A. Rincon, J. Correa-Bordes, J.C. Ribas, and P. Perez. 2018. Paxillin-Mediated Recruitment of Calcineurin to the Contractile Ring Is Required for the Correct Progression of Cytokinesis in Fission Yeast. Cell reports. 25:772–783 e774.

22. McCollum, D., A. Feoktistova, and K.L. Gould. 1999. Phosphorylation of the myosin-II light chain does not regulate the timing of cytokinesis in fission yeast. The Journal of biological chemistry. 274:17691–17695.

23. Mochizuki, T., G. Wu, T. Hayashi, S.L. Xenophontos, B. Veldhuisen, J.J. Saris, D.M. Reynolds, Y. Cai, P.A. Gabow, A. Pierides, W.J. Kimberling, M.H. Breuning, C.C. Deltas, D.J. Peters, and S. Somlo. 1996. PKD2, a gene for polycystic kidney disease that encodes an integral membrane protein. *Science (New York*, N.Y). 272:1339–1342.

24. Moreno, S., A. Klar, and P. Nurse. 1991. Molecular genetic analysis of fission yeast Schizosaccharomyces pombe. Methods in enzymology. 194:795–823.

25. Morris, Z., D. Sinha, A. Poddar, B. Morris, and Q. Chen. 2019. Fission yeast TRP channel Pkd2p localizes to the cleavage furrow and regulates cell separation during cytokinesis. Molecular biology of the cell. 30:1791–1804.

26. Naqvi, N.I., K.C. Wong, X. Tang, and M.K. Balasubramanian. 2000. Type II myosin regulatory light chain relieves auto-inhibition of myosin-heavy-chain function. Nature cell biology. 2:855–858.

27. Palmer, C.P., E. Aydar, and M.B. Djamgoz. 2005. A microbial TRP-like polycystic-kidney-disease-related ion channel gene. The Biochemical journal. 387:211–219.

28. Petronczki, M., P. Lenart, and J.M. Peters. 2008. Polo on the Rise-from Mitotic Entry to Cytokinesis with Plk1. Developmental cell. 14:646–659.

29. Poddar, A., Y.Y. Hsu, F. Zhang, A. Shamma, Z. Kreais, C. Muller, M. Malla, A. Ray, A. Liu, and Q. Chen. 2022. Membrane stretching activates calcium-permeability of a putative channel Pkd2 during fission yeast cytokinesis. Molecular biology of the cell:mbcE22070248.

30. Poddar, A., O. Sidibe, A. Ray, and Q. Chen. 2021. Calcium spikes accompany cleavage furrow ingression and cell separation during fission yeast cytokinesis. Molecular biology of the cell. 32:15–27.

31. Pollard, T.D., and J.Q. Wu. 2010. Understanding cytokinesis: lessons from fission yeast. Nature reviews. 11:149–155.

32. Prieto-Ruiz, F., E. Gomez-Gil, R. Martin-Garcia, A.J. Perez-Diaz, J. Vicente-Soler, A. Franco, T. Soto, P. Perez, M. Madrid, and J. Cansado. 2023. Myosin II regulatory light chain phosphorylation and formin availability modulate cytokinesis upon changes in carbohydrate metabolism. eLife. 12.

33. Sinha, D., D. Ivan, E. Gibbs, M. Chetluru, J. Goss, and Q. Chen. 2022. Fission yeast polycystin Pkd2p promotes cell size expansion and antagonizes the Hippo-related SIN pathway. Journal of cell science. 135.

34. Stark, B.C., T.E. Sladewski, L.W. Pollard, and M. Lord. 2010. Tropomyosin and myosin-II cellular levels promote actomyosin ring assembly in fission yeast. Molecular biology of the cell. 21:989–1000.

35. Thevenaz, P., U.E. Ruttimann, and M. Unser. 1998. A pyramid approach to subpixel registration based on intensity. IEEE Trans Image Process. 7:27–41.

36. Wu, J.Q., J.R. Kuhn, D.R. Kovar, and T.D. Pollard. 2003. Spatial and temporal pathway for assembly and constriction of the contractile ring in fission yeast cytokinesis. Developmental cell. 5:723–734.

37. Wu, J.Q., and T.D. Pollard. 2005. Counting cytokinesis proteins globally and locally in fission yeast. *Science (New York*, N.Y). 310:310–314.

38. Yoshida, T., T. Toda, and M. Yanagida. 1994. A calcineurin-like gene ppb1+ in fission yeast: mutant defects in cytokinesis, cell polarity, mating and spindle pole body positioning. Journal of cell science. 107 (Pt 7):1725–1735.

